# Cell-Projection Pumping of Fibroblast Contents into Osteosarcoma SAOS-2 Cells Correlates with Increased SAOS-2 Proliferation and Migration, and also with Altered Morphology

**DOI:** 10.1101/2021.11.23.469761

**Authors:** Swarna Mahadevan, James A Cornwell, Belal Chami, Elizabeth Kelly, Hans Zoellner

**Author notes:** (SM); (EK).

## Abstract

We earlier reported that cell-projection pumping transfers fibroblast contents to cancer cells, and this alters cancer cell phenotype. We now report on single-cell tracking of time lapse recordings from co-cultured fluorescent fibroblasts and SAOS-2 osteosarcoma cells, tracking 5,201 cells across 7 experiments. The fluorescent lipophilic marker DiD was used to label fibroblast organelles, and to trace transfer of fibroblast cytoplasm into SAOS-2. We related SAOS-2 phenotypic change to levels of fluorescence transfer from fibroblasts to SAOS-2, and also to what we term ‘compensated fluorescence’, that numerically projects mother cell fluorescence post-mitosis, into daughter cells. Comparison of absolute with compensated fluorescence, allowed deduction if phenotypic effects in mother SAOS-2, were inherited by their daughters. SAOS-2 receipt of fibroblast fluorescence correlated by Kendall’s tau: with cell-profile area, and without evidence for persistence in daughter cells (median tau = 0.51, *p* < 0. 016); negatively and weakly with cell circularity, and with evidence for persistence (median tau = −0.19, *p* < 0.05); and very weakly with cell migration velocity, and without evidence for persistence (median tau = 0.01, *p* < 0.016). Also, mitotic SAOS-2 had higher rates of prior fluorescence uptake (median = 64.9 units/day), compared with non dividing cells (median = 35.6 units/day, *p* < 0.016), and there was no evidence for persistence post-mitosis. We conclude there is appreciable impact of cell-projection pumping on cancer cell phenotype, relevant to cancer histopathological diagnosis, clinical spread, and growth, with most effects ‘reset’ by cancer cell mitosis.

## 1. Introduction

We earlier described the exchange of membrane, organelles and cytoplasmic protein between cultured human fibroblasts and cancer cells [1–3], and others have made similar observations [4–27]. Uptake of cellular contents by such transfers may be via exosomes, tunnelling nanotubes, or a mechanism we recently reported and term ‘cell-projection pumping’, and these can significantly change the phenotype of acceptor cells [2–20]. Transfer of mitochondria appears especially significant, and can confer chemotherapy resistance to cancer cells [4, 6, 7, 20–27]. However, we are made wary of ascribing most phenotypic effects to mitochondrial exchange, by our observation that in addition to mitochondria, there is also bulk transfer of: cytoplasmic proteins; plasma membrane bound alkaline phosphatase; and organelles smaller than mitochondria [1–3].

Exosomes comprise membrane bound vesicular structures shed by cells, that are readily taken up by neighboring cells [11–13, 17, 18]. Tunnelling nanotubes, are delicate tube-like structures that establish cytoplasmic continuity between often distant cells, and in two-dimensional cell cultures appear as pipe-like structures suspended above the culture surface [8–10, 14, 15, 28–30]. Cell-projection pumping is a hydrodynamic mechanism, whereby retracting cell-projections inject their cytoplasmic contents into adjacent cells [3]. A recent independent report supports the novel cell-projection pumping mechanism, and relates chemotherapy resistance of multiple myeloma cells, to uptake of mitochondria from co-cultured bone marrow derived stromal cells [21].

Our time-lapse microscopy observations show that cell-projection pumping is the main mechanism for transfer of fluorescently labelled human dermal fibroblast (HDF) contents, into co-cultured SAOS-2 osteosarcoma cells. More modest transfers are also seen from SAOS-2 into HDF. Mathematical modelling and computer simulation demonstrates this mechanism can account for all observable fluorescence transfers in this culture system [3]. For this reason, it is convenient to study co-cultures of HDF with SAOS-2 for purposes of exploring the biological significance of cell-projection pumping.

Morphological analysis of SAOS-2 in co-cultures with human gingival fibroblasts, reveals reduced cell circularity and increased cell-profile area in SAOS-2 that have received fluorescent fibroblast label [1]. When SAOS-2 are separated from co-cultured fibroblasts by transwell membranes, fibroblast cytokine synthesis is markedly different to when cell-projection pumping is permitted by direct contact [31]. Furthermore, SAOS-2 co-cultured with fluorescently labelled HDF, can be separated by fluorescence activated cell sorting (FACS) into those with high and low levels of HDF label uptake [2]. Co-cultured SAOS-2 with high levels of HDF label and separated from their fellows by FACS, have increased cell migration in scratch assays [2]. Also, FACS analysis demonstrates that SAOS-2 with high levels of HDF fluorescent label, have increased cell size and increased internal structural complexity, and both these changes are consistent with the notion that cell-projection pumping transfers fibroblast organelles and cytoplasm into SAOS-2 [2].

Emergence of cancer cell diversity is considered a critical factor for cancer progression and resistance to treatment [13, 32]. While outgrowth of genetically distinct sub-clones of cancer cells is a key driver for cancer cell diversity, interactions of cancer cells with stromal cells in the surrounding tumour microenvironment are increasingly thought important [4, 6, 7, 13, 20, 21, 32–35]. Considered in this light, our data exploring SAOS-2 phenotypic changes following receipt of fibroblast contents [1–3, 31], suggest that cell-projection pumping may contribute to clinically important cancer cell diversity and the tumour microenvironment, where: SAOS-2 morphological changes [1, 2], are relevant to histopathological diagnosis [36]; altered cytokine synthesis [31], contributes to the tumour microenvironment [33, 35]; and increased SAOS-2 migration [2], contributes to cancer spread through the body [32, 37]. Although we have focused on fibroblast co-cultures with SAOS-2, we have observed similar transfers and phenotypic changes, when fibroblasts are co-cultured with other cancer cell lines including from: melanomas; colon carcinomas; ovarian carcinomas; lung cancer and osteosarcomas [1, 2].

While our published work is informative on cell-projection pumping and some aspects of the possible significance for cancer, we have been considering limitations of the methods we used. One major difficulty, is that changes in SAOS-2 phenotype are related to levels of fibroblast fluorescence uptake, but any fibroblast label a given SAOS-2 may have received, is effectively halved when the cell divides. The effect of this on FACS separated SAOS-2 populations, is to shift the daughter cells of SAOS-2 that may have had high levels of fibroblast label, into the ‘low fibroblast label’ population, thus undermining assays. Although results for cell proliferation in FACS separated co-cultured SAOS-2 were negative, it occurred to us that increased SAOS-2 proliferation could have been masked by such ‘fluorescence halving’, and we felt unable to make clear conclusion if cell-projection pumping affected SAOS-2 cell division [2]. Also, we see that our earlier morphological analysis of co-cultured SAOS-2 in fixed monolayers [1], would have been similarly affected by SAOS-2 division, and also provided no information on the discrete history of individual cells.

We now address these limitations by single-cell tracking of SAOS-2 co-cultured with HDF. This method records the behavior and fate of individual cells and their progeny across multiple cell divisions and generations [38–40]. By identifying the fate of individual cells, single-cell tracking overcomes the limitations of pooled cell assays that average the outcomes for thousands of cells. An interesting finding of single-cell tracking studies, is that sister cells are more similar to each other, than they are to their mother cell or their own progeny [38–40].

It was especially interesting for us to explore the possibility that the memory of HDF cytoplasm received by SAOS-2 mother cells, might be preserved in daughter cells. In this study, we assess the effect of cell-projection pumping from HDF into SAOS-2 on: cell-profile area; cell circularity; cell migration velocity; and proliferation. We relate SAOS-2 phenotype to absolute levels of fluorescence acquired from HDF, as well as to what we term ‘compensated fluorescence’ that assigns to each paired daughter cell, half of their mother cell’s fluorescence which is lost to each sister on mother cell division. Taking the experience of mother cells into account in this way, and comparing results for absolute with compensated florescence, we were able to deduce if phenotypic effects were inherited by their daughters, or if mitosis reset SAOS-2 to their native state. We further relate divergence in the phenotype of paired sister cells, to differences in levels of HDF contents received.

## 2. Materials and Methods

### 2.1. Materials

All cell culture media, phosphate buffered saline (PBS), trypsin (0.25%)/EDTA (1mM) and bovine calf serum (BCS) were purchased from Thermo Fisher Scientific (Waltham, MA, USA). Gelatin was from Sigma-Aldrich (St. Louis, MO, USA). Tissue culture plasticware was purchased from Costar (Cambridge, MA, USA). CSL Biosciences (Parkville, VIC, Australia) supplied antibiotics penicillin and streptomycin. ICN Biomedicals Inc. (Costa Mesa, CA, USA) provided amphotericin B. HDF were from The Coriell Institute (Camden, NJ, USA). SAOS-2 osteosarcoma cells were from the American Type Culture Collection (Manassas, VA, USA). The lipophilic fluorescent probes DiD (excitation 644nm, emission 665nm) and DiO (excitation 484nm, emission 501nm) Vybrant cell labelling solutions were from Molecular Probes, Life Technologies (Grand Island, NY, USA). 24 well black-bottomed plates were from Ibidi (Gräfelfing, Barvaria, Germany).

### 2.2. Cell Culture

The antibiotics penicillin (100 U/mL), streptomycin (100 μg/mL) and amphotericin B (2.5μg/mL) were used throughout all cell culture. HDF were always cultured on gelatin coated surfaces (0.1% in PBS) in DMEM (15% BCS). SAOS-2 were cultured in DMEM with BCS (10%). Cells were harvested using trypsin-EDTA, into BCS to neutralize trypsin, and pelleted by centrifugation before passage at a ratio of 1 to 3. All cell culture was performed at 37° C under CO_2_ (5%) and at 100% humidity.

### 2.3. Labelling of Cells with Lipophilic Fluorescent Membrane Markers

Labelling solutions of DiD (1mM) and DiO (2mM) were prepared in DMEM with 10% BCS, and applied to cells for 1 h. Cells were then washed twice with PBS before overnight culture with DMEM with BCS (15%), followed by two further washes with PBS in order to ensure removal of any unbound label [1–3].

### 2.4. Co-Culture Conditions

All experiments were performed with cells cultured in gelatin coated (0.1% in PBS) black bottomed 24 well culture plates. HDF were seeded at from 1 to 2 x 10^4^ cells per cm^2^ and allowed to adhere overnight before labelling with DiD and further overnight culture in DMEM with BCS (2%) as outlined above. SAOS-2 were seeded prior to labelling at near confluence in M199 with BCS (15%), and allowed to adhere overnight before labelling with DiO and further overnight culture in DMEM with BCS (2%) as outlined above. Pre-labelled SAOS-2 were then seeded over HDF in DMEM with BCS (10%) at a culture density of 4 x 10^4^ cells per cm^2^ for up to 5 days co-culture, with experiments terminated as monolayers approached confluence. Seven separate experiments were conducted, and coded ‘a’ to ‘g’. One experiment was conducted over 2 days (experiment b), 1 over 3.6 days (experiment c), four over 4 days (experiments a, d, e and g), and one over 5 days (experiment f). Media were changed at day 3 for experiments extending 4 and 5 days. Control cultures comprised HDF and SAOS-2 labelled and seeded in parallel for isolated cell culture.

### 2.5. Time-lapse recordings

Experimental culture plates were placed in a humidified culture chamber that was mounted on a Leica DM 16000B fluorescence and phase contrast microscope, and maintained under CO_2_ (5%) at 37° C. Viewing cells through a 20x objective, Micro-Manager open source software [41] was used to construct 3 x 3 grids of contiguous visual fields, each measuring 643 μm x 482 μm. Slight overlap of adjacent visual fields was made to avoid potentially disruptive gaps, so that grids measuring approximately 1,920 μm x 1,440 μm were observed in each well. Recordings were made of fluorescently labelled co-cultured cells, as well as of fluorescently labelled HDF and SAOS-2 control cells cultured alone. Recording of images was controlled by Micro-Manager software, using a Leica DFC365 FX camera. Phase contrast images were collected at 15 min intervals. Fluorescence images were acquired at a lower rate of one image per 4 hr, to minimize photobleaching and phototoxicity. Final recordings were assembled into coherent time-lapse sequences using code developed in MATLAB (MathWorks, Natick, MA, USA).

### 2.6. Single-cell tracking and analysis

Single-cell tracking software was adapted from that previously developed in MATLAB (TrackPad: https://github.com/Jamcor/TrackPad), and was used to follow the fate of individual cells and their progeny in phase contrast time-lapse recordings [38, 39]. Cells present at the beginning of experiments, were defined as the ‘starting population’, while sequential cell divisions generated ‘first’, ‘second’, ‘third’ and very occasionally ‘fourth’ generations of progeny. The number of cells tracked in each experiment is given for co-cultures of HDF and SAOS-2 (Table S1, Supplementary Materials) and in control cells cultured in isolation (Table S2, Supplementary Materials). Across all experiments: 1,846 co-cultured SAOS-2 were tracked from 607 starting cells; 992 co-cultured HDF were tracked from 523 starting cells; 1,514 control SAOS-2 were tracked from 540 starting cells; and 849 control HDF were tracked from 458 starting cells. From this, a total of 5,201 cells were tracked in the current study. Details of how precise progeny relationships were coded are given in the ‘Explanatory Notes’ for Supplementary Materials, S2, being an Excel spreadsheet with all experimental data summarised.

The ultimate fate of all tracked cells was determined as either: cell division; incomplete division; apoptosis; or ‘incomplete’ meaning the cell was either lost from view or reached the end of the experiment. Discrete progeny relationships between all tracked cells were unambiguous. Individual cells were segmented for analysis of cell circularity and cell-profile area only at times that both phase contrast and fluorescence images were available, that is only at multiples of 4 hourly intervals. Where a tracked cell either: divided, underwent apoptosis; failed to divide; or was lost to the field of vision, its phase contrast image was segmented at the time of the immediately prior fluorescence images. The cell position in the x-y position of the image was recorded, as well as the time of: phase contrast and fluorescence image capture; cell appearance on mother cell mitosis; cell apoptosis; loss from view of the cell; or mitosis. Because the mitotic arrival of cells in the starting generation was undefined, it was not possible to determine intermitotic time or the time till apoptosis, for starting generation cells.

### 2.7. Segmentation of cells and dependent calculations for cell circularity, absolute fluorescence and compensated fluorescence

Cells were manually segmented from phase contrast images, giving results for cell-profile area and cell peripheral circumference. Cell circularity was calculated from these by the equation: cell circularity = 4pi(Cell profile area) / (cell peripheral circumference)^2.^. DiD fluorescence was quantitated for segmented cells, by summation of intensity of red pixels in fluorescence images, masked by the shape of the segmented cells.

Data was imported into RStudio open source software [42]. We were interested to compare the relationship of SAOS-2 cell phenotype with levels of DiD fluorescence accepted from co-cultured HDF. We were further interested to consider if phenotypic effects were carried post-mitosis into later generations of cells. To explore this possibility, SAOS-2 cell phenotype was related to two separate measures for receipt of HDF fluorescence. Absolute fluorescence of cells (Fa) comprised the DiD fluorescence observed in SAOS-2. Compensated fluorescence from the mother cell (Fmc) was calculated for tracked cells that arose by mitosis during the experiment. This awarded to each of the two sister cells, half of the mother cell’s fluorescence, thus compensating for distribution of mother cell fluorescence to both daughters. It was assumed that cell division distributed mother cell fluorescence equally to both daughters, so that the numeric correction comprised addition to Fa of each daughter, of half of the immediate mother cell’s Fa value. Separate preliminary analyses compensating for fluorescence from all ancestor generations of cells was also attempted, but it was realized that variance inherent to measurements made, exceeded the levels of accuracy needed for those calculations to have meaning.

### 2.8. Determination of cell migration velocity

Moving cell location was determined during single-cell tracking in Trackpad, while further analysis was in RStudio, similar to earlier reports [38, 39]. In brief, mean cell migration velocity was calculated for all tracked cells, by firstly identifying cell centroids in phase contrast images at relevant time points at 2 h intervals, performed in Trackpad. Distances migrated between centroids and time points were calculated in RStudio as in a cartesian plot, that is by the square root of the summated squares of differences for horizontal and vertical coordinates. All distances that a given cell had travelled were then summated, and divided by the total time the cell was tracked. Please note that the 2h interval was selected on basis that distances travelled for shorter time intervals were often within the range of error for microscope stage relocation. The last time point frequently did not coincide with the 2 h interval, and where the last time interval was less than 2 h, the preceding time point was removed to create a final time interval greater than 2h.

### 2.9. Normalization of fluorescence values

To aid comparison across experiments, despite variability inherent to DiD fluorescence labelling, Fa and Fmc values for all SAOS-2 were normalized relative to the median DiD fluorescence of co-cultured SAOS-2 in the starting generation, which was defined as having a value of 100 normalized fluorescence units.

### 2.10. Correlation by Kendall’s tau of cell-profile area, cell circularity and cell migration velocity with receipt of HDF fluorescence

Cell-profile area, cell circularity and migration velocity of individual SAOS-2 cells were compared separately against both Fa and Fmc. Within individual experiments, correlation was assessed by Kendall’s tau, and statistical significance of this was noted. Kendall’s tau was determined for individual generations of cells within experiments, as well as for sequentially added generations to both summarize results, and examine possible confounding effects from prolonged culture or cell crowding.

### 2.11. Correlation of mitosis with receipt of HDF fluorescence

Cells were identified as either undergoing mitosis or not, and because this was ‘statistically nominal data’, it was not possible to use Kendall’s tau to examine association between receipt of fluorescence and SAOS-2 cell division. Instead, receipt of fluorescence by SAOS-2 that subsequently underwent division, was directly compared with that in cells that did not experience mitosis. It was inherent to the culture system, that mitosis was asynchronous and that the times that dividing and non dividing cells arose varied greatly, as did the times for which these cells were observed. To account for this, values for Fa and Fmc were divided by the time of observation, so that it was the rate of fluorescence uptake (Fa/day and Fmc/day) that was examined, when considering effects of receipt of HDF fluorescence on SAOS-2 division.

### 2.12. Calculation of an index for persistence of phenotypic effect inherited from mother cells

An ‘index of persistence’ was calculated to assess inheritance by daughter cells, of any phenotypic effects of fluorescence uptake observed in their respective mitotic mother cells. In this index, a value of ‘0’ indicated no evidence for persistence; and ‘1’ indicated strong evidence for persistence of the effect. Please note that the calculated persistence index did not relate to the strength of persistence of phenotype into descendent cells. Instead, the index related to the proportionate number of experiments where evidence of persistence was seen.

The rationale for calculation of the persistence index for cell-profile area, cell circularity and cell migration velocity, was that if there was persistence of effect beyond mitosis, disorder would be introduced to any correlation of Fa with the phenotypic effect, thus reducing the strength of Kendall’s tau. Numerical compensation of fluorescence in Fmc would, however, improve order and so increase the strength of Kendall’s tau. On the other hand, in absence of persistence of effect beyond mitosis, numerical compensation of fluorescence would introduce disorder for correlation with Fmc, reducing the strength of Kendall’s tau relative to that for Fa. From this, by comparing Fa with Fmc, it was possible to deduce whether or not the effect of receiving HDF fluorescence on the phenotype of mother cells, survived mitosis to be inherited by the two daughter cells. For individual experiments, a value of +1 was assigned where tau for Fa < tau for Fmc; 0 was assigned where tau for Fa = tau for Fmc; and −1 was assigned where tau for Fa > tau for Fmc. Where negative correlations were involved, all tau values were multiplied by −1 to maintain comparability of final results. Results of these comparisons for all experiments were then summated and averaged, to yield scores ranging from −1 to 1. Final persistence index values ranging from 0 to 1 were calculated from these averaged scores as per: persistence index = (1 + Final Score)/2.

Because data on mitosis was unique amongst other recordings made, in being ‘statistically nominal’, a modified approach was required to calculate the persistence index for the effect of receiving HDF fluorescence on SAOS-2 division. Values for Fa and Fmc in dividing cells, were first divided by those in non dividing cells, to yield ‘proportionate fluorescence values’ pFa and pFmc for each experiment. A higher pFmc relative to pFa, was interpreted as evidence for persistence with assignation of a score of +1. Lower or equivalent pFmc relative to pFa was interpreted as evidence against persistence of effect, with assignation of a score of −1 for the individual experiment. Further summation, averaging and final calculation of persistence index, was as described above for other phenotypic features.

### 2.13. Analysis of the relationship between acquired fluorescence and phenotype in paired SAOS-2 sister cells

It is logical to expect that where there is clear effect on SAOS-2 phenotype of uptake of HDF fluorescence, that there would be correlation when the differences between paired sister cells in phenotype, are plotted against differences in fluorescence acquired by the paired sister cells. Sister cells by definition can only be identified from generation 1 onwards, and Kendall’s tau was determined for differences between paired sister cells in: cell-profile area; cell circularity; cell migration velocity; and inter-mitotic time.

### 2.14. Evaluation of statistical significance

Statistical significance was accepted for *p* < 0.05. Statistical significance of divergence of Kendall’s tau from an expected correlation of 0 was evaluated by the One Sample Wilcoxon Test. Statistical significance of differences in Kendall’s tau between Fa and Fmc for all experiments, as well as of differences in phenotype between paired groups of cells, was by the Wilcoxon Signed Rank Test. The Mann Whitney U Test was used to compare unpaired results. While most statistical evaluations were performed in RStudio, it was occasionally convenient to perform analyses using Prism software (9.2.0, Graphpad Software, San Diego, CA, USA).

## 3. Results

### 3.1. There was marked transfer of HDF fluorescent label to SAOS-2 during co-culture

HDF and SAOS-2 labelled clearly with DiD and DiO fluorescent markers, and when these cells were co-cultured there was appreciable and highly localized transfer of DiD from HDF to SAOS-2, typical of that expected for cell-projection pumping (Figure 1).

**Figure 1.**
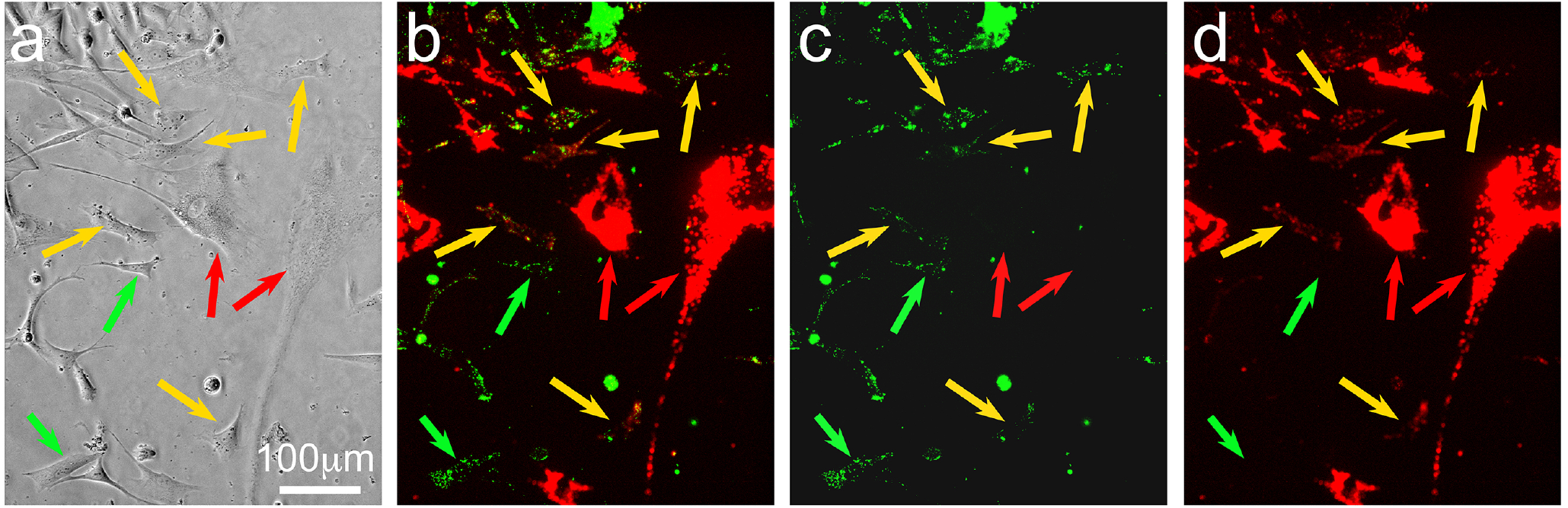
Photomicrographs of a visual field of pre-labelled SAOS-2 (fluorescent green marker) co-cultured with pre-labelled HDF (fluorescent red marker) at the 20h time point, showing phase contrast (a) and fluorescent images with red and green channels combined (b), as well as for green (c) and red (d) channels alone. The lipophilic fluorescent markers were concentrated in organelles, creating a typically punctate appearance (b,c,d), so that phase contrast images (a) were more helpful for identification of cell margins. HDF were clearly labelled red (red arrows), while SAOS-2 were labelled green. Comparison of combined and separated channels for fluorescence images, demonstrates that some SAOS-2 had no red HDF fluorescent label (green arrows), while others had received appreciable levels of the red HDF marker (yellow arrows).

### 3.2. SAOS-2 had lower cell-profile area and migration velocity, and higher cell circularity compared with HDF

Although the cell-profile area of control SAOS-2 and HDF cultured in isolation varied both within and between experiments, SAOS-2 were consistently different compared with HDF (Table S3, Supplementary Materials). The median value of cell-profile area in SAOS-2 cultured in isolation was 1,359 μm^2^, which was significantly lower than that in HDF in isolated cell culture (median value 4,172 μm^2^), while cell circularity in SAOS-2 was higher than in HDF (SAOS-2 median of 0.70; HDF median of 0.36, *p* < 0.016; Wilcoxon Signed Rank Test). Separately, HDF had significantly faster cell migration than SAOS-2 (HDF median of 288 μm/day; SAOS-2 median of 84 μm/day; *p* < 0.016, Wilcoxon Signed Rank Test). These general differences between SAOS-2 and HDF in cell-profile area, circularity and migration velocity, were retained when the cells were co-cultured (*p* < 0.016, Wilcoxon Signed Rank Test).

### 3.3. SAOS-2 cell-profile area correlated with receipt of HDF fluorescence and the effect did not persist post-mitosis

Figure 2 shows typical results, in this case from experiment ‘a’, with strong correlation between cell-profile area in co-cultured SAOS-2 and both Fa and Fmc, and in this experiment for all generations of cells tracked. In addition to considering individual generations of cells, it was convenient to summarize effects by examining correlation when successive generations of cells were analyzed together. As shown in Figure 2 for example, starting and first generations were considered as a single group to yield one pair of Kendall’s tau values for Fa and Fmc, while a still further pair of tau values were obtained when all generations were considered together.

**Figure 2.**
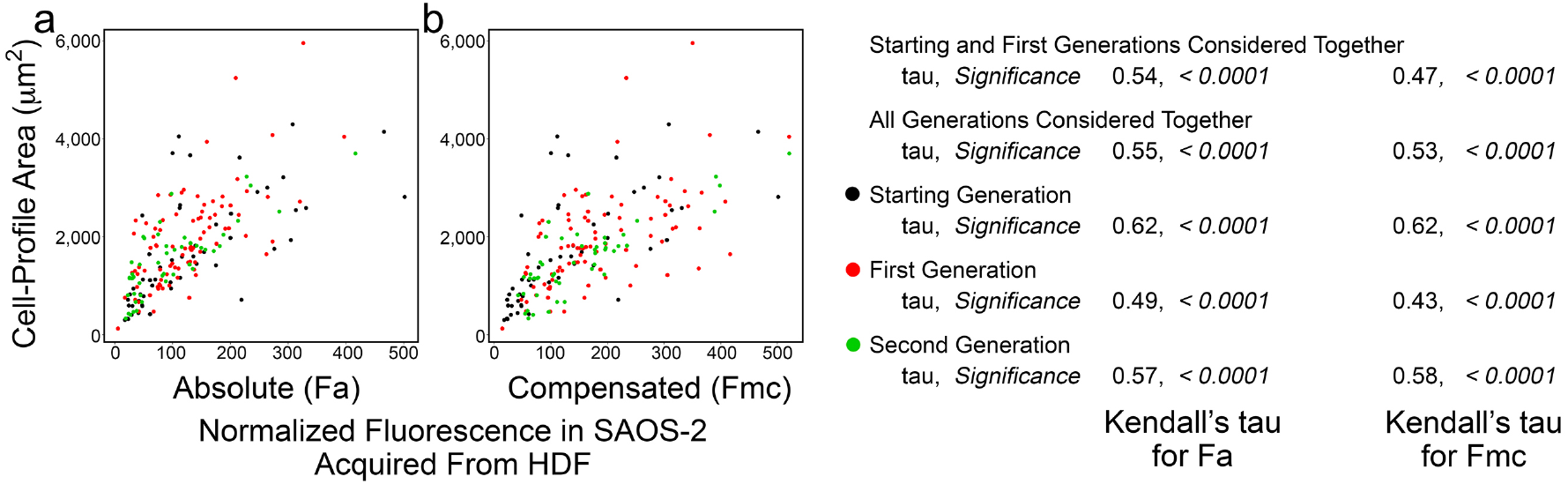
Scatter plots of data from experiment ‘a’ co-culturing SAOS-2 with HDF, showing cell-profile area of co-cultured SAOS-2 plotted against normalized fluorescence acquired from HDF expressed as absolute fluorescence measured (Fa in panel a), as well as with numerical compensation for halving of fluorescence by cell division in mother cells (Fmc in panel b). The generation to which each cell belonged is indicated by color. Values for Kendall’s tau of correlation and statistical significance of this are shown for: individual generations of cells; starting and first generations together; and all generations of cells together. (a) There was correlation between cell-profile area and Fa. (b) This reduced for Fmc.

Results were similar for all experiments (Table 1), and when cells of all generations were considered together, the correlation between receipt of HDF fluorescence and SAOS-2 cell-profile area was stronger for Fa than for Fmc (*p* < 0.016, Wilcoxon Signed Rank Test). This was generally retained when cells across earlier sequential generations were considered (Supplementary Materials, Table S4). The reduced correlation in Fmc relative to Fa was visually reflected in scatter plots, by an increasing spread of data points (Figure 2). There was no evidence for persistence of this effect on phenotype in daughter cells, after mother cell division (Table 1). There was correlation between differences in Fa of paired sister cells, and difference in cell-profile area of these cells (Figure 3, Table S5 in Supplementary Materials, *p* < 0.032 One Sample Wilcoxon Test considering generations 1 and 2 together).

**Table 1.**
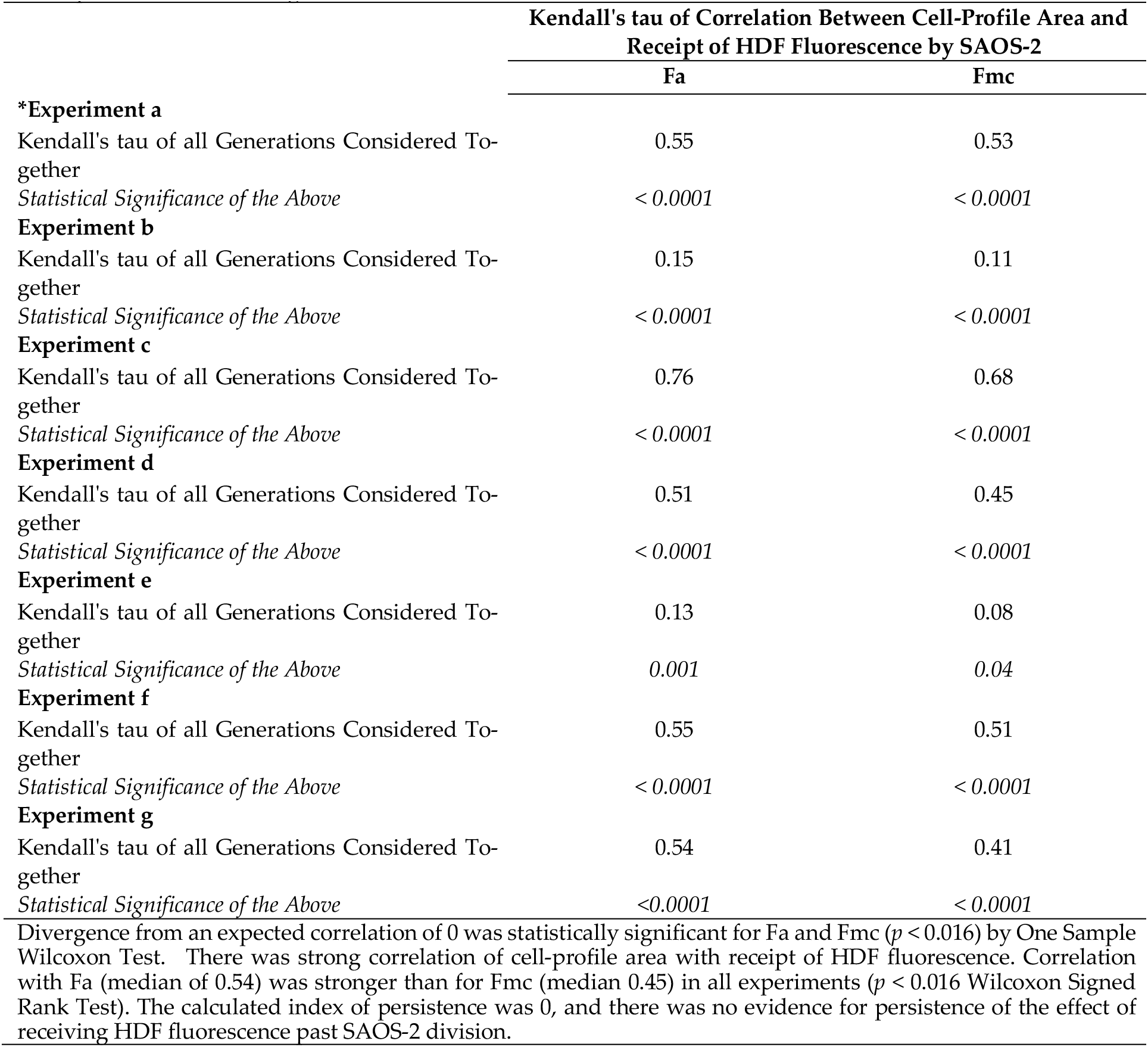
Kendall’s tau of correlation between cell-profile area of tracked SAOS-2 and absolute fluorescence acquired from co-cultured HDF (Fa), as well as with compensation for halving of fluorescence from mother cells by cell division (Fmc). Results for all experiments are shown, considering all generations of cells counted together. Results for other groupings of successive generations of cells are in Supplementary Materials Table S4. * Indicates the experiment shown in Figure 2.

|  | Kendall's tau of Correlation Between Cell-Profile Area and Receipt of HDF Fluorescence by SAOS-2 |  |
| --- | --- | --- |
|  | Fa | Fmc |
| **\*Experiment a** |  |  |
| Kendall's tau of all Generations Considered Together | 0.55 | 0.53 |
| *Statistical Significance of the Above* | $< 0.0001$ | $< 0.0001$ |
| **Experiment b** |  |  |
| Kendall's tau of all Generations Considered Together | 0.15 | 0.11 |
| *Statistical Significance of the Above* | $< 0.0001$ | $< 0.0001$ |
| **Experiment c** |  |  |
| Kendall's tau of all Generations Considered Together | 0.76 | 0.68 |
| *Statistical Significance of the Above* | $< 0.0001$ | $< 0.0001$ |
| **Experiment d** |  |  |
| Kendall's tau of all Generations Considered Together | 0.51 | 0.45 |
| *Statistical Significance of the Above* | $< 0.0001$ | $< 0.0001$ |
| **Experiment e** |  |  |
| Kendall's tau of all Generations Considered Together | 0.13 | 0.08 |
| *Statistical Significance of the Above* | 0.001 | 0.04 |
| **Experiment f** |  |  |
| Kendall's tau of all Generations Considered Together | 0.55 | 0.51 |
| *Statistical Significance of the Above* | $< 0.0001$ | $< 0.0001$ |
| **Experiment g** |  |  |
| Kendall's tau of all Generations Considered Together | 0.54 | 0.41 |
| *Statistical Significance of the Above* | $< 0.0001$ | $< 0.0001$ |
| Divergence from an expected correlation of 0 was statistically significant for Fa and Fmc ($p < 0.016$) by One Sample Wilcoxon Test. There was strong correlation of cell-profile area with receipt of HDF fluorescence. Correlation with Fa (median of 0.54) was stronger than for Fmc (median 0.45) in all experiments ($p < 0.016$ Wilcoxon Signed Rank Test). The calculated index of persistence was 0, and there was no evidence for persistence of the effect of receiving HDF fluorescence past SAOS-2 division. | | |

**Figure 3.**
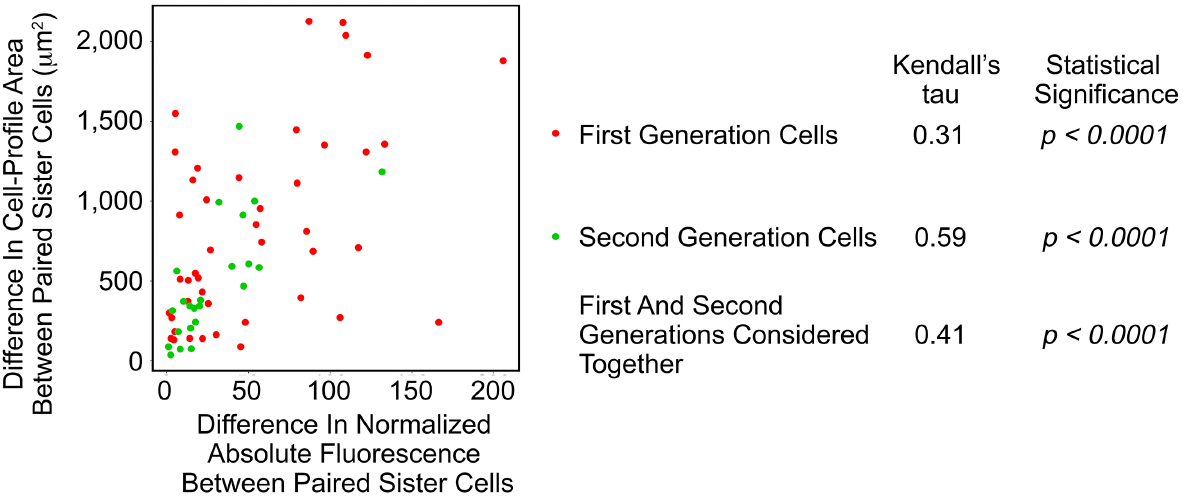
Scatterplot of the differences between paired sister SAOS-2 cells in cell-profile area against differences in fluorescence acquired from co-cultured HDF. Data are from the same experiment ‘a’, which is shown in Figure 2. There was correlation in both generations of cells studied, as well as when first and second generatinos were considered together.

### 3.4. Cell circularity in co-cultured SAOS-2 was inversely correlated with receipt of HDF fluorescence and the effect persisted post-mitosis

Figure 4 shows results from experiment ‘a’ where there was weak although statistically significant inverse correlation between Fa and cell circularity in co-cultured SAOS-2. The inverse correlation was also seen between cell circularity and Fmc. This correlation was weaker when all generations were considered together (Figure 4), as was generally the case across most experiments (Supplementary Materials Table S6), and this likely reflected confounding effects of cell crowding at later time points. For this reason, analysis across experiments was of results for the starting and first generation grouped (Table 2).

**Table 2.**
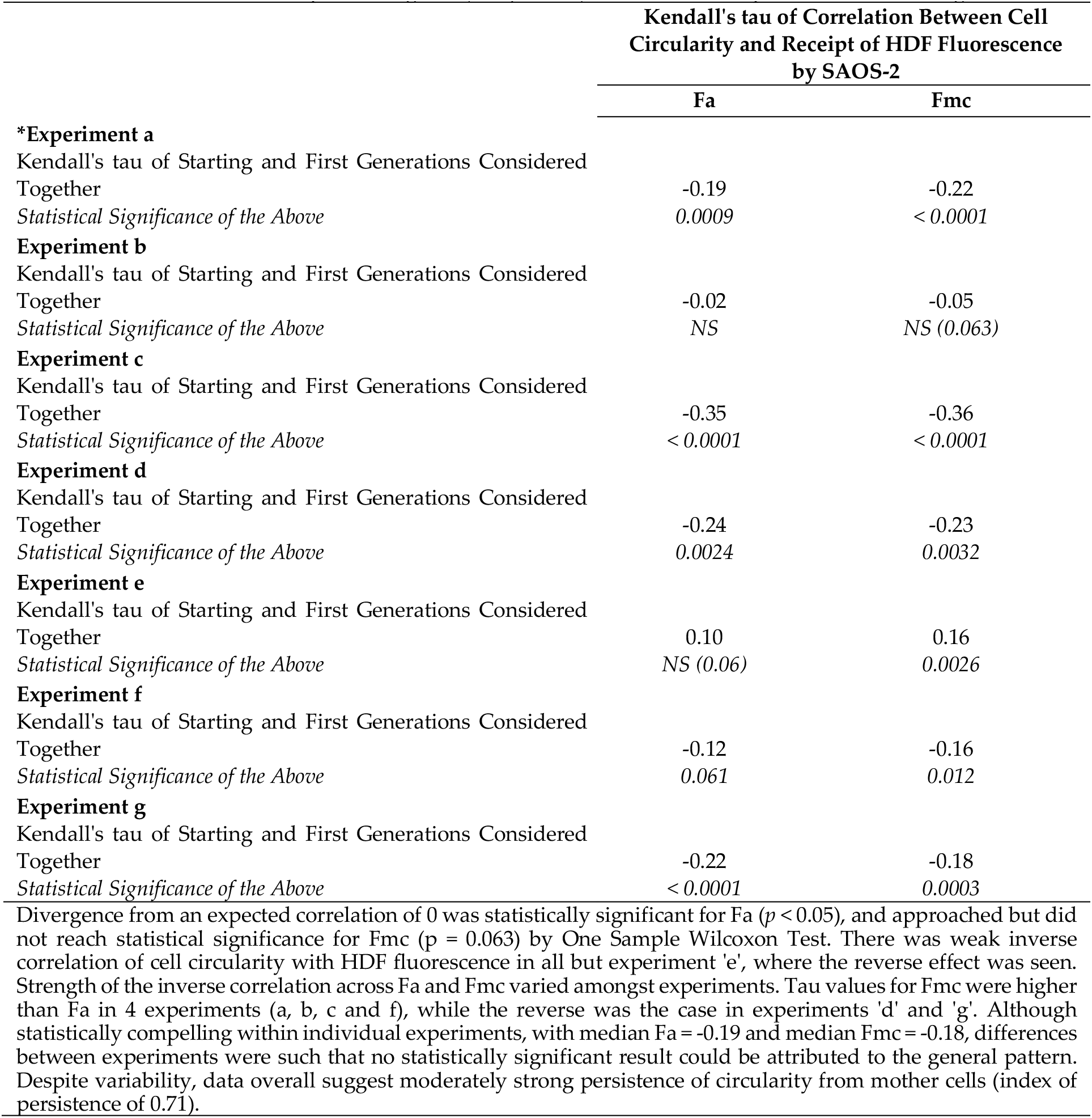
Kendall’s tau of correlation between cell circularity of tracked SAOS-2 and absolute fluorescence acquired from co-cultured HDF (Fa), as well as with compensation for halving of fluorescence from mother cells by cell division (Fmc). Results for all experiments are shown, considering starting and first generations together. Results for other groupings of successive generations of cells are in Supplementary Materials Table S6. Statistical significance is given, where *NS* indicates ‘not significant’ to *p* < 0.05. Where statistical significance was approached but not reached, the calculated p value is given (*NS (p value)*). *Indicates the experiment shown in Figure 4.

**Figure 4.**
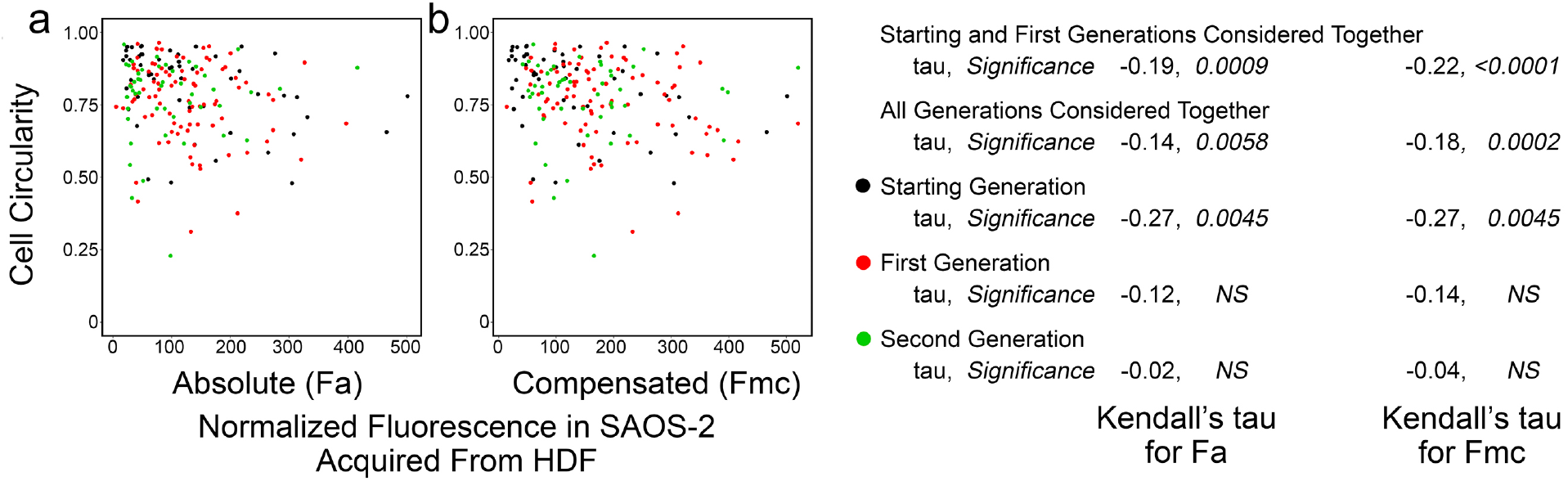
Scatter plots of experiment ‘a’ co-culturing SAOS-2 with HDF, showing cell circularity of co-cultured SAOS-2 plotted against normalized fluorescence acquired from HDF expressed as absolute fluorescence measured (Fa in panel a), as well as with numerical compensation for halving of fluorescence by cell division from mother cells (Fmc in panel b). The generation to which each cell belonged is indicated by color. Values for Kendall’s tau of correlation and statistical significance of this are shown for: individual generations of cells; starting and first generations together; and all generations of cells together. (a) There was inverse correlation between cell-profile and Fa. (b) This increased for Fmc.

The strength and statistical significance of the inverse correlation was higher for Fmc than for Fa in four experiments (Table 2, Supplementary Materials Table S6). The reverse, however, was seen in two experiments (Table 2, Supplementary Materials Table S6), while in one experiment the correlation seemed reversed. Modest persistence of the effect past mother cell division was apparent from the calculated persistence index of 0.71. Table S7 (Supplementary Materials) shows that it was in only two experiments (e and g) that any correlation was seen in differences between sister cells for cell circularity and acquired fluorescence. In keeping with the correlations between Fa and Fmc with cell circularity, this correlation too was weak, and was only statistically significant when cells from all generations were considered together (experiment e, Kendall’s tau = 0.12, *p* < 0.05; experiment f, Kendall’s tau = 0.14, *p* < 0.035).

### 3.5. SAOS-2 migration velocity had weak correlation with receipt of HDF fluorescence and the effect did not persist post-SAOS-2 division

Weak correlation was seen between Fa and cell migration velocity, but this did not reach statistical significance in some experiments (Figure 5, Table 3, Supplementary Materials Table S8). Similar correlation was seen for Fmc, but this was less often statistically significant and occasionally reversed (Table 3). There was negligible evidence for persistence beyond cell division of this modest phenotypic effect (Table 3). Also, there was no correlation for differences between paired sister cells in migration velocity and fluorescence uptake (Table S9, Supplementary Materials).

**Table 3.**
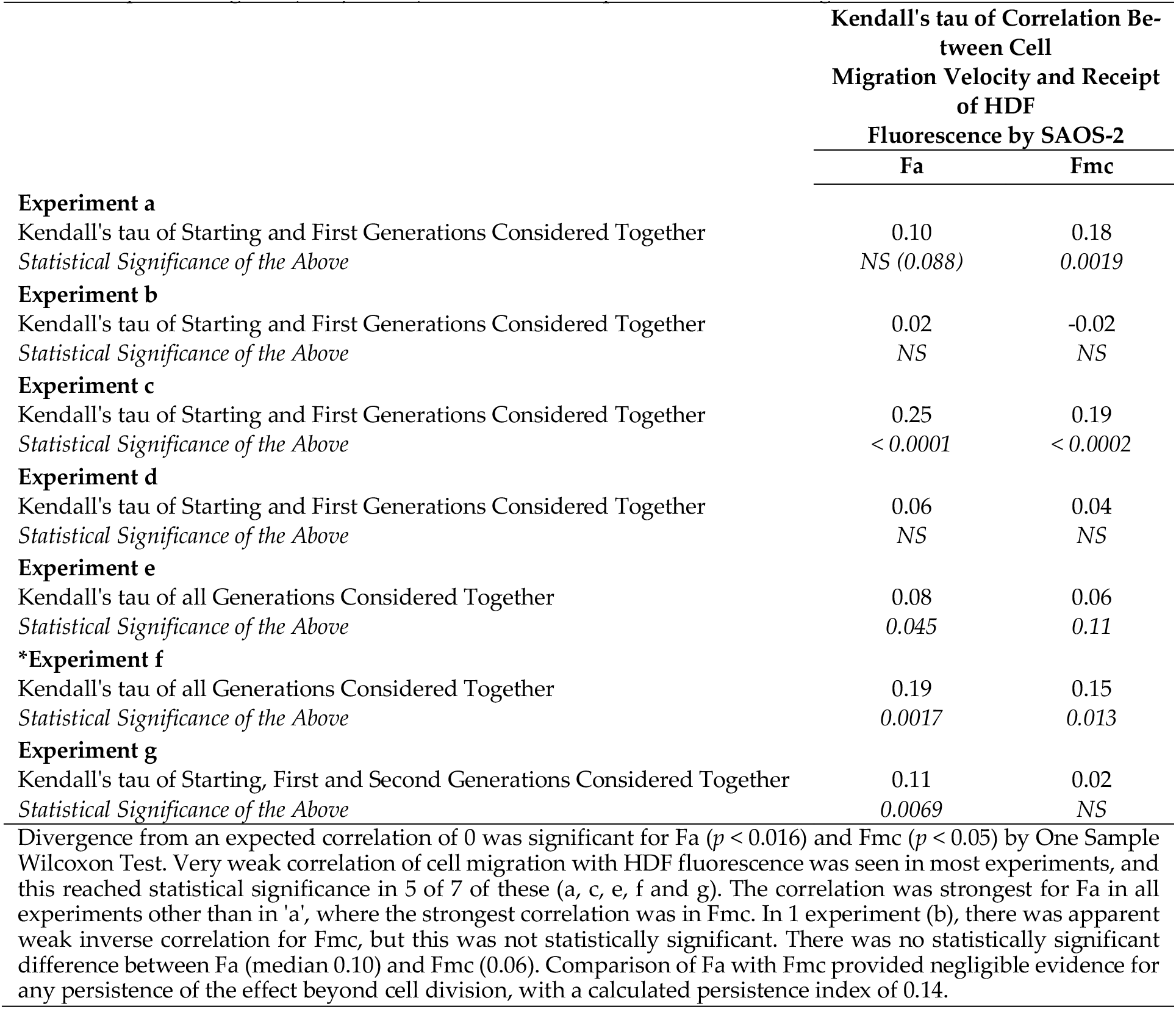
Kendall’s tau of correlation between cell migration velocity of tracked SAOS-2 and absolute fluorescence acquired from co-cultured HDF (Fa), as well as with compensation for halving of fluorescence from mother cells by cell division (Fmc). Results from all experiments are shown. Results shown within individual experiments are from either: all generations of cells together; starting and first generations together; or where a second generation of cells was present, starting, first and second generations together. Selection of these within experiments was on basis of the strongest statistical significance, while results for all grouped cell generations are shown in Supplementary Materials Table S8. Note that there were no statistically significant differences in experiments b and d, and selection here was from starting and first generations considered together. Statistical significance is given, where *NS* indicates ‘not significant’ to *p* < 0.05. Where statistical significance was approached but not reached, the calculated p value is given (*NS (p value)*). *Indicates the experiment shown in Figure 5.

|  | Kendall's tau of Correlation Between Cell Migration Velocity and Receipt of HDF Fluorescence by SAOS-2 |  |
| --- | --- | --- |
|  | Fa | Fmc |
| **Experiment a** |  |  |
| Kendall's tau of Starting and First Generations Considered Together | 0.10 | 0.18 |
| *Statistical Significance of the Above* | NS (0.088) | 0.0019 |
| **Experiment b** |  |  |
| Kendall's tau of Starting and First Generations Considered Together | 0.02 | -0.02 |
| *Statistical Significance of the Above* | NS | NS |
| **Experiment c** |  |  |
| Kendall's tau of Starting and First Generations Considered Together | 0.25 | 0.19 |
| *Statistical Significance of the Above* | < 0.0001 | < 0.0002 |
| **Experiment d** |  |  |
| Kendall's tau of Starting and First Generations Considered Together | 0.06 | 0.04 |
| *Statistical Significance of the Above* | NS | NS |
| **Experiment e** |  |  |
| Kendall's tau of all Generations Considered Together | 0.08 | 0.06 |
| *Statistical Significance of the Above* | 0.045 | 0.11 |
| **\*Experiment f** |  |  |
| Kendall's tau of all Generations Considered Together | 0.19 | 0.15 |
| *Statistical Significance of the Above* | 0.0017 | 0.013 |
| **Experiment g** |  |  |
| Kendall's tau of Starting, First and Second Generations Considered Together | 0.11 | 0.02 |
| *Statistical Significance of the Above* | 0.0069 | NS |
| Divergence from an expected correlation of 0 was significant for Fa ($p < 0.016$) and Fmc ($p < 0.05$) by One Sample Wilcoxon Test. Very weak correlation of cell migration with HDF fluorescence was seen in most experiments, and this reached statistical significance in 5 of 7 of these (a, c, e, f and g). The correlation was strongest for Fa in all experiments other than in 'a', where the strongest correlation was in Fmc. In 1 experiment (b), there was apparent weak inverse correlation for Fmc, but this was not statistically significant. There was no statistically significant difference between Fa (median 0.10) and Fmc (0.06). Comparison of Fa with Fmc provided negligible evidence for any persistence of the effect beyond cell division, with a calculated persistence index of 0.14. | | |

**Figure 5.**
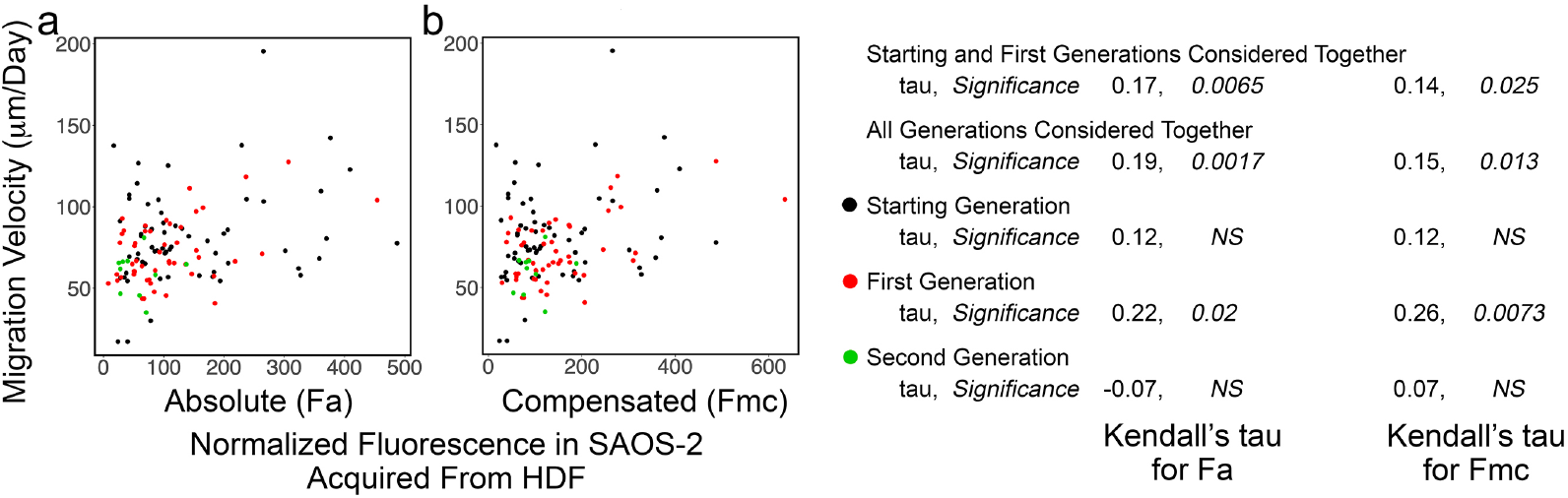
Scatter plots of experiment ‘f’ co-culturing SAOS-2 with HDF, showing cell migration velocity of co-cultured SAOS-2 plotted against normalized fluorescence acquired from HDF expressed as absolute fluorescence measured (Fa in panel a), as well as with numerical compensation for halving of fluorescence by cell division in mother cells (Fmc in panel b). The generation to which each cell belonged is indicated by color. Values for Kendall’s tau of correlation and statistical significance of this are shown for: individual generations of cells; starting and first generations together; and all generations of cells together. (a) There was weak correlation between cell migration and Fa. (b) This decreased for Fmc.

### 3.6. Increased HDF fluorescence transfer to SAOS-2 during co-culture was associated with subsequent SAOS-2 mitosis and there was no evidence for persistence of this post-cell division

Mitosis was frequently seen amongst tracked cells (622 of 1,846 co-cultured SAOS-2; 235 of 992 co-cultured HDF; 488 of 1,514 control SAOS-2 cultured in isolation; 198 of 849 control HDF cultured in isolation), to produce sequential generations of cells (Tables S1 and S2, Supplementary Materials). Apoptosis was much less common (114 of 1,846 co-cultured SAOS-2; 18 of 992 co-cultured HDF; 98 of 1,514 control SAOS-2 cultured in isolation; 13 of 849 control HDF cultured in isolation), and did not appear to contribute significantly results (Tables S1 and S2, Supplementary Materials).

To evaluate the effect of receiving HDF fluorescence on SAOS-2 mitosis, co-cultured SAOS-2 from all generations were considered together, excluding late generations where there was either insufficient experimental time for cell division to occur, or where there were an insufficient number of dividing and or non dividing cells for proper comparison. Cells that subsequently underwent division, had generally higher rates of both Fa and Fmc uptake than those that did not (Figure 6, Table 4). There was negligible evidence for persistence of the effect post-mitosis (Table 4).

**Table 4.** The fate of SAOS-2 with regard to cell division, related to the rate of previous fluorescence uptake from co-cultured HDF. Median rates of acquisition of fluorescence are given for absolute fluorescence acquired per day (Fa/day), and fluorescence compensating for halving on division of immediate mother cells (Fmc/day). Statistical significance (Stat. Sig.) as per the Mann Whitney U Test is given, where *NS* indicates ‘not significant’ to *p* < 0.05. Where statistical significance was approached but not reached, the calculated p value is shown (*NS (p value)*). All generations of cells for which adequate data was available were included for analysis. However, in experiment ‘a’, first generation cells had atypical results compared with all other experiments with disproportionate effect on pooled results. For this reason, two results for experiment ‘a’ are shown, one including these cells, and another where only cells in the starting generation of experiment ‘a’ were analysed (Experiment a*). Both pFa and pFmc to assess persistence of effect are shown for all experiments other than for experiment ‘a*’ where pFmc has no meaning.

|  | Median Fa / day |  |  | Median Fmc / day |  |  |
| --- | --- | --- | --- | --- | --- | --- |
|  | Dividing Cells | Non Dividing Cells | pFa<br>(Dividing Cells /<br>Non Dividing Cells) | Dividing Cells | Non Dividing Cells | pFmc<br>(Dividing Cells /<br>Non Dividing Cells) |
| Experiment a | 49.0 | 76.9 | 0.64 | 61.1 | 120.1 | 0.51 |
| Stat. Sig. of Above | 0.0022 |  |  | < 0.0001 |  |  |
| Experiment a* | 45.6 | 35.6 | 1.28 | 45.6 | 35.6 | - |
| Stat. Sig. of Above | NS |  |  | NS |  |  |
| Experiment b | 89.4 | 73.3 | 1.22 | 113.5 | 153.3 | 0.74 |
| Stat. Sig. of Above | NS (0.057) |  |  | NS |  |  |
| Experiment c | 66.7 | 29.9 | 2.23 | 51.4 | 11.4 | 4.51 |
| Stat. Sig. of Above | < 0.0001 |  |  | < 0.0001 |  |  |
| Experiment d | 75.0 | 65.8 | 1.88 | 102.0 | 100.1 | 1.02 |
| Stat. Sig. of Above | NS (0.0535) |  |  | NS |  |  |
| Experiment e | 39.9 | 15.8 | 2.53 | 73.0 | 50.0 | 1.46 |
| Stat. Sig. of Above | < 0.0001 |  |  | 0.0041 |  |  |
| Experiment f | 45.6 | 32.6 | 1.40 | 49.1 | 39.2 | 1.25 |
| Stat. Sig. of Above | 0.0222 |  |  | NS |  |  |
| Experiment g | 64.9 | 42.1 | 1.54 | 94.0 | 65.9 | 1.43 |
| Stat. Sig. of Above | 0.0005 |  |  | 0.0115 |  |  |
| Co-cultured SAOS-2 that underwent cell division had higher rates of uptake for Fa, than SAOS-2 that did not divide (Excluding experiment 'a': median Fa/day for dividing cells = 65.8; median Fa/day for non dividing cells = 37.3; $p < 0.032$, Wilcoxon Signed Rank Test. Including experiment a*: median Fa/day for dividing cells = 64.9; median Fa/day for non dividing cells = 35.6; $p < 0.016$, Wilcoxon Signed Rank Test). This was also seen in 5 of 6 relevant experiments for Fmc (c, d, e, f and g), although there was negligible difference in experiment 'd', and the reverse was seen in experiment 'b' (median Fmc/day for dividing cells = 73.0; median Fmc/day for non dividing cells = 50.0, not statistically significant). Excluding experiment 'a', comparison of pFa with pFmc yielded a negligibly low persistence index of 0.17, so that persistence of the association between uptake of fluorescence and cell division in descendent cells was not supported by these data. | | | | | | |

**Figure 6.**
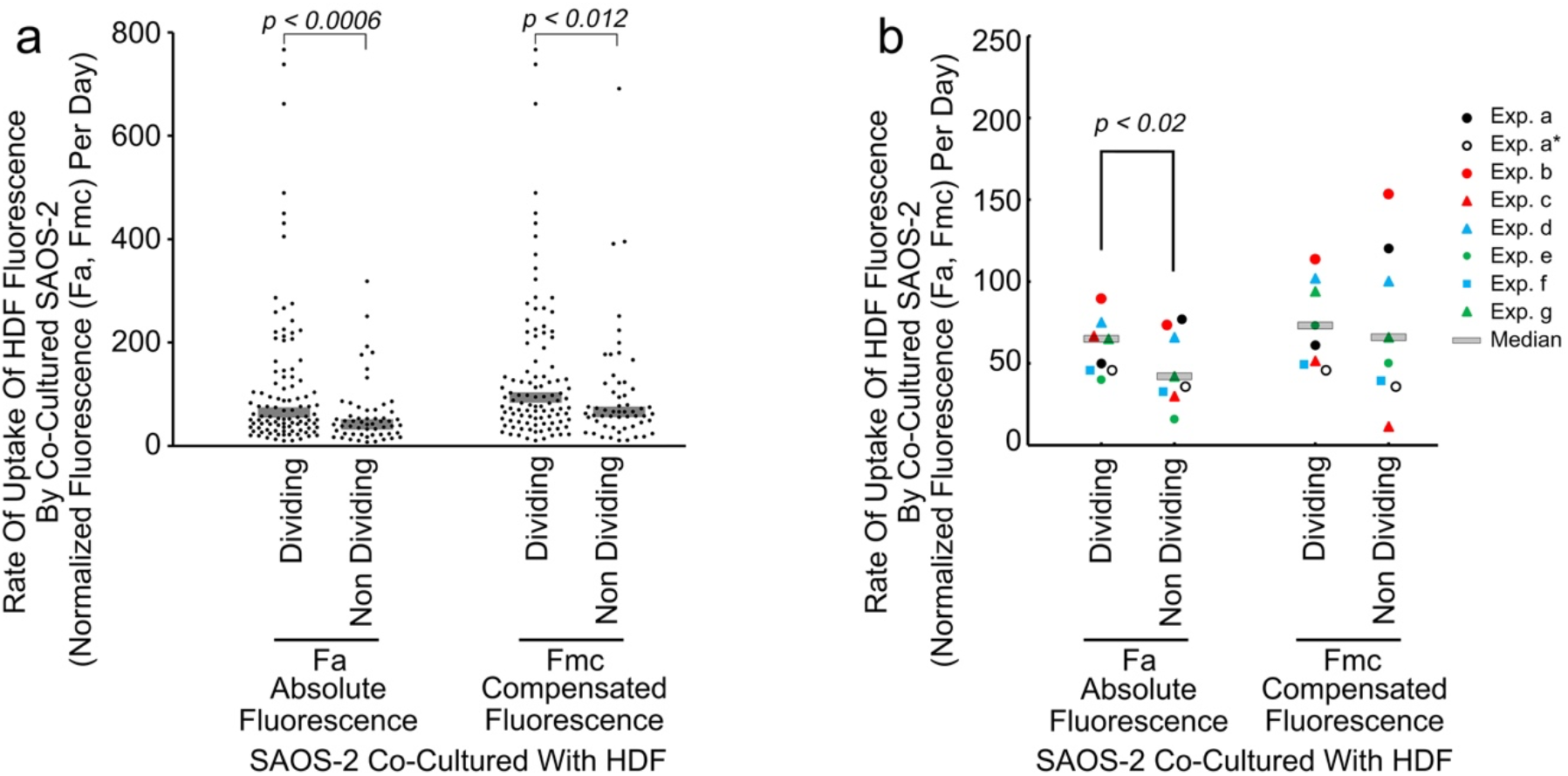
Scattergram showing results of a typical experiment (experiment g) showing the rate of uptake of normalized HDF fluorescence by individual co-cultured SAOS-2 cells, according to whether cells underwent subsequent cell division or not (a), as well as median values from all experiments (b). Fluorescence is expressed as the rate of uptake of absolute fluorescence measured (Fa/day), as well as with numerical compensation for halving of fluorescence by cell division in mother cells (Fmc/day). (a) Dividing cells had clearly higher values for Fa/day than did non dividing cells, and this was statistically significant as marked (Mann Whitney U Test). A similar statistically significant difference was seen for Fmc/day. (b) Results for all experiments are shown, while experiment ‘a’ which showed atypical results for first generation cells is indicated twice, once including first generation cells, and once without (Exp. a*). The median rate of Fa uptake was higher in those cells that subsequently underwent division than in those that did not, and this was statistically significant as marked when experiment ‘a’ first generation cells only were considered, and also when experiment ‘a’ was excluded from analysis (*p* < 0.032, Wilcoxon Signed Rank Test). The relationship for Fmc was similar although less consistent, and was not statistically significant when all experiments were considered together.

## 4. Discussion

Findings of the current work are broadly consistent with the work of others who report increased cell migration and or proliferation following transfer of cell contents via tunneling nanotubes [7, 22–26], as well as with our own earlier reports [1, 2]. Although special interest has been shown in the literature for the role of mitochondria [4, 6, 7, 20–27], our observation of bulk cytoplasmic transfer via cell-projection pumping, including of cytoplasmic protein, plasma membrane alkaline phosphatase and organelles smaller than mitochondria [1–3], makes us cautious of focusing on this single organelle as the critical factor for all changes observed. Instead, we suggest that in addition to mitochondria, any cellular component transferred, has potential to profoundly affect the recipient cell.

One reason for using single-cell tracking in the current study, was interest in exploring whether or not the phenotypic impact of fibroblast transfers to cancer cells, transcended mitosis to affect daughter cells. A particular challenge to addressing this question, was that post mitotic SAOS-2 were open to receive still further transfers from HDF, obscuring any phenotypic effects inherited from mother cells. It seems reasonable for us to have assumed that receipt of HDF fluorescence was a good proxy for the amount of HDF contents received and phenotypic effect in mother cells, and that there was equal distribution of mother cell fluorescence to each mitotic daughter cell. On that basis, our approach of comparing correlation of phenotype against Fa, with that for Fmc, to deduce persistence or otherwise of phenotypic effects post-mitosis, also seems reasonable. At the level of individual experiments, it was possible to make direct comparison of Kendall’s tau for Fa and Fmc for cell-profile area, cell circularity and cell migration velocity, and of pFa with pFmc with regard to cell division. There was, however, some variability in outcomes of experiments, so in addition to evaluation with statistical tests appropriate to include multiple experiments, it was helpful to determine an index of persistence. We find no clear precedent for our approach in the literature, and suggest it may find application elsewhere by others.

It seemed very clear, that associations between receipt of HDF fluorescence and SAOS-2 cell-profile area, migration velocity and mitosis, were reset by cell division. This suggests that whatever it is that HDF cell-projection pumping delivers to SAOS-2 that drives these phenotypic changes, it is either degraded or otherwise unstable, so that the effects do not survive mitotic events. This is consistent with separate reports of the importance of scheduled protein and RNA degradation, associated with cell cycle progression, suggested by others as important for resetting cells after mitosis [43–51]. Also, because mitochondrial function is retained post-mitosis [52], the current findings reinforce our doubt of the possible importance of mitochondria, in accounting for the changes we describe.

With regard to cell circularity, however, a degree of persistence was observed. It might be thought that the modest persistence observed was artifactual due to the low levels of correlation involved. This would, however, be inconsistent with absence of persistence for cell migration velocity, where correlation was even weaker. Persistence of circularity effects post-mitosis could reflect epigenetic changes in SAOS-2, but given variability across experiments, it more likely that there is degradation of the driver or drivers responsible for cell circularity similar to those for other phenotypic effects studied, but that these are sufficiently robust to penetrate post-mitosis into daughter cells. Clarity awaits characterization of the precise contents transferred from HDF, and of the mechanisms responsible for effecting these phenotypes.

Separately, we exploited the high similarity of sister cells [38–40], to help verify phenotypic effects of fluorescence transfers seen. It was reassuring to see the expected correlation in this for cell-profile area. Although divergence in cell-circularity for sister cells was only seen in two experiments, this was perhaps to be expected given the much weaker level of correlation involved. Similar arguments apply to the absence of correlation between paired sister SAOS-2 with regard to cell migration velocity, where the direct correlation with Fa and Fmc was even weaker than for cell circularity.

A further motivation for using single-cell tracking, was to overcome potential confounding effects of pooled cell assays in earlier work [1, 2]. In this regard, outcomes of the current study were mixed. Of benefit was that the current study for the first time, associated receipt of fibroblast label with SAOS-2 mitosis. We earlier used FACS to separate SAOS-2 with high levels of HDF fluorescence uptake from those with low uptake. In that work, we found no proliferative effect [2], which at first light seems inconsistent with the current findings by single-cell tracking. Since, however, we demonstrate in addition to the pro-mitotic association, failure of this to persist in daughter cells, we see that these two studies can be reconciled considering the distribution through the FACS column of daughter cells with high levels of inherited mother cell fluorescence, and the pooled cell proliferation assay that extended over several days and cell divisions [2]. It appears that mitosis can be driven while SAOS-2 have opportunity to receive new transfers from fibroblasts during actual co-culture, but that once isolated from the fibroblasts as for example by FACS separation, the cell-projection pumping driver for mitosis is lost.

The single-cell approach had further benefit over our initial study of fixed monolayers of cells, where increased cell-profile area and reduced cell circularity were seen, but only at the cell population level as shifts in data for SAOS-2 that had clear fluorescence uptake, relative to those that did not [1]. The current demonstration of changes in cell circularity and cell-profile area at the single cell level, increases confidence in our earlier findings [1]. This is especially the case for cell-profile area, where data from paired sister cells was particularly convincing. We previously demonstrated by FACS analysis, that uptake of HDF contents by SAOS-2 correlates with increased size of SAOS-2 [2], and it is tempting to imagine the current correlation of fibroblast fluorescence transfer with increased SAOS-2 cell-profile area, is a simple expression of that. However, cell-profile area in two dimensional culture is more reflective of active cell stretching than of mere volume, so we suspect this observation indicates a change in SAOS-2 cell activity, rather than passive cell size. In as much as the current single-cell tracking study had advantages over previous pooled cell assays [1, 2], there were also some disadvantages. We found that data were sensitive to culture density effects, with for example data on cell circularity and cell migration velocity from late generations of cells, being affected by increased cell crowding.

We were particularly surprised to find only slight correlation between receipt of HDF fluorescence and SAOS-2 migration velocity, because in our earlier work with FACS isolated cells, the effect was marked [2]. We suggest that in addition to stimulating SAOS-2 migration via cell-projection pumping, HDF are also able to inhibit SAOS-2 migration in co-culture. This possibility is supported by the work of others, who report that fibroblasts can inhibit cancer cell migration by both soluble and contact dependent mechanisms [53]. Further surprising to us, was absence of evidence for persistence of increased migration post-mitosis, because our earlier work with FACS isolated cells in scratch assays, showed increased migration over many days and clearly over multiple cell divisions [2]. It is possible that there might be persistence of effect, but that that the low levels of correlation observed in the current study, make that undetectable by the methods used. Another possibility, is that the leading edge of migrating cells in scratch assays, comprised starting generation cells that did not divide but effectively filtered themselves out of the pooled population, by migrating into scratches. Histopathological cancer grading systems often include assessment of variability in morphology amongst cancer cells, formally described as pleomorphism, with high pleomorphism usually indicating higher grade and worse clinical outcomes [32]. The current findings on cell-profile area and cell circularity are consistent with our earlier suggestion that cell-projection pumping increases morphological cancer cell diversity, and that this has bearing on cancer diagnosis [1].

While significance of the effect of HDF cell-projection pumping on SAOS-2 migration now seems complicated by what may be opposing effects, as outlined above, the significance of the association with increased mitosis and hence cancer growth, does seem clinically important.

Our interest in persistence or otherwise of the effects on SAOS-2 of receiving HDF contents, and demonstration that most effects are reset by mitosis, could distort perception of the clinical significance of cell-projection pumping. It thus seems important to note, that *in-vivo*, fibroblasts are nearly always available to provide fresh contents to newly arrived cancer cells, so that irrespective of persistence, cancer cells are always open to the impact of cell-projection pumping, which on basis of our data, we believe to be important. Since the cell-projection pumping mechanism is only recently described [3], we envision future development of novel anti-cancer therapies that target cell-projection pumping.

## 5. Conclusions

We conclude that transfer of HDF contents to SAOS-2 via cell-projection pumping, drives: increased SAOS-2 mitosis; modestly increased SAOS-2 migration; increased SAOS-2 cell profile area; and reduced SAOS-2 circularity. In addition, we conclude that all these phenotypic changes, other than cell circularity, are reset by SAOS-2 mitosis. We suggest this indicates an important role for cell-projection pumping in clinical cancer histopathologic diagnosis, and progression of the disease. We envision development of novel anti-cancer therapies targeting cell-projection pumping.

## Supporting information

Excel File with Data

## Supplementary Materials

The following are available online at www.mdpi.com/xxx/s1: S1. Supplementary Tables, Table S1. The number of cells tracked in co-cultures of SAOS-2 with HDF, according to cell type, generation and experiment; Table S2. The number of control SAOS-2 and HDF tracked, according to cell type, generation and experiment; Table S3. Median values for cell-profile area, cell circularity and cell migration velocity, in control SAOS-2 and HDF cultured in isolation for all experiments. Table S4 Kendall’s tau of correlation between cell-profile area of tracked SAOS-2 and Fa and Fmc; Table S5. Kendall’s tau for correlation in differences between paired sister cells for cell-profile area and Fa; Table S6. Kendall’s tau of correlation between cell circularity of tracked SAOS-2 and Fa and Fmc; Table S7. Kendall’s tau for correlation of differences between paired sister cells for cell circularity and Fa; Table S8. Kendall’s tau of correlation between cell migration velocity of tracked SAOS-2 and Fa and Fmc; Table S9. Kendall’s tau for correlation of differences between paired sister cells for cell migration velocity and Fa. S2. Supplementary Experimental Data. A zipped Excel file with spreadsheets containing key data for all cells studied, and explanatory notes on the coding system used to identify progeny relationships between cells.

## Author Contributions

SM and HZ conceived experiments, conducted numerical and statistical analyses, and prepared the first drafts of the manuscript. Further, SM performed experiments, and conducted both single-cell tracking and downstream RStudio analysis. JC played a leading role establishing and tailoring single-cell tracking software for the project and in early analysis. EK assisted with cell culture. BC supported microscopy. All authors contributed to manuscript preparation. The authors declare no conflict of interest.

## Funding

We thank the Australian Dental Research Fund for their support of this work. We also thank an anonymous donor for their kind contribution.

## Data Availability Statement

Data for re-analysis of results are provided in with Supplementary Materials.

## Acknowledgments

In addition to funders of this work, we wish to thank Dr R Nordon of the Graduate School of Biomedical Engineering, University of NSW for initial development of Trackpad software, and for his generosity sharing this.

## Conflicts of Interest

The corresponding author (HZ) is the Director and Head of Strongarch Pty Ltd, which is a freelance academic company. Strongarch and HZ have no commercial conflict of interest in this work, and the remaining authors also declare no conflicts of interest

## Notes

### Competing Interest Statement

The authors have declared no competing interest.

