## Supplementary Tables for "Cell-Projection Pumping of Fibroblast Contents into Osteosarcoma SAOS-2 Cells Correlates with Increased SAOS-2 Proliferation and Migration, and also with Altered Morphology"

[1] The Cellular and Molecular Pathology Research Unit, Oral Pathology and Oral Medicine, School of Dentistry, Faculty of Medicine and Health, The University of Sydney, Westmead Hospital, Westmead, NSW, 2145, Australia; (SM); (EK)
[2] Laboratory of Cancer Biology and Genetics, Center for Cancer Research, National Cancer Institute, Bethesda, MD, 20892, USA;
[3] Molecular Biomedicine, Charles Perkins Centre, The University of Sydney, NSW, 2006, Australia;
[4] Biomedical Engineering, Faculty of Engineering, The University of Sydney, NSW, 2006, Australia
[5] Graduate School of Biomedical Engineering, University of NSW, NSW, 2052, Australia
[6] Strongarch Pty Ltd, Sydney, NSW, 2000, Australia

### S1. Supplementary Tables

**Table S1.** The number of cells tracked in co-cultures of SAOS-2 with HDF, according to cell type, generation (Gen.), experiment, and apoptosis, indicated by numbers in brackets (n).

| | The Number of Co-Cultured SAOS-2 Tracked | | | | | |
|---|---|---|---|---|---|---|
| | Starting Gen. | 1st Gen. | 2nd Gen. | 3rd Gen. | 4th Gen. | Total |
| Experiment a | 52 | 88 (1) | 46 (1) | | | 186 (2) |
| Experiment b | 234 (19) | 312 (16) | 86 | | | 632 (35) |
| Experiment c | 98 (10) | 81 (13) | 4 | | | 183 (23) |
| Experiment d | 33 (4) | 44 (2) | 40 (1) | 8 | | 125 (7) |
| Experiment e | 51 | 99 (6) | 121 (4) | 31 (1) | | 302 (11) |
| Experiment f | 63 (6) | 50 (8) | 10 | | | 123 (14) |
| Experiment g | 76 (14) | 112 (6) | 103 (2) | 4 | | 295 (22) |
| Total | 607 (53) | 786 (52) | 410 (8) | 43 (1) | | 1,846 (114) |
| | The Number of Co-Cultured HDF Tracked | | | | | |
| | Starting Gen. | 1st Gen. | 2nd Gen. | 3rd Gen. | 4th Gen. | Total |
| Experiment a | 37 | 30 | 16 | 6 | | 89 |
| Experiment b | 168 (4) | 120 (3) | 26 (1) | 4 | | 318 (8) |
| Experiment c | 105 (4) | 28 (1) | 10 (1) | | | 143 (6) |
| Experiment d | 31 (1) | 33 | 32 (1) | 4 | | 100 (2) |
| Experiment e | 70 | 66 (1) | 42(1) | 18 | 2 | 198 (2) |
| Experiment f | 53 | | | | | 53 |
| Experiment g | 59 | 24 | 8 | | | 91 |
| Total | 523 (9) | 301 (5) | 134 (4) | 32 | 2 | 992 (18) |

Considering all co-cultured cells across all 7 experiments, a total of 2,838 co-cultured cells were tracked, amongst which 132 became apoptotic. The number of mitotic events is clear from the number of cells tracked in successive generations. Note that very occasionally, a daughter cell was quickly lost from vision and so not tracked, accounting for odd numbers in columns for generations 1 onwards, that contain otherwise paired sister cells.

**Table S2.** The number of control SAOS-2 and HDF tracked, according to cell type, generation (Gen.), experiment, and apoptosis, indicated by numbers in brackets (n).

| | The Number of Control SAOS-2 Tracked | | | | | |
|---|---|---|---|---|---|---|
| | Starting Gen. | 1st Gen. | 2nd Gen. | 3rd Gen. | 4th Gen. | Total |
| Experiment a | 46 (6) | 62 | 14 | | | 122 (6) |
| Experiment b | 147 (10) | 176 (9) | 17 | | | 340 (19) |
| Experiment c | 129 (8) | 154 (15) | 23 (1) | | | 306 (24) |
| Experiment d | 38 (3) | 56 (3) | 54 (6) | 6 | | 154 (12) |
| Experiment e | 88 (4) | 146 (10) | 120 (4) | 14 | 2 | 370 (18) |
| Experiment f | 50 (8) | 30 (2) | | | | 80 (10) |
| Experiment g | 42 | 66 (9) | 34 | | | 142 (9) |
| Total | 540 (39) | 690 (48) | 262 (11) | 20 | 2 | 1,514 (98) |

| | The Number of Control HDF Tracked | | | | | |
|---|---|---|---|---|---|---|
| | Starting Gen. | 1st Gen. | 2nd Gen. | 3rd Gen. | 4th Gen. | Total |
| Experiment a | 37 | 39 | 24 | 3 | 2 | 105 |
| Experiment b | 78 (2) | 31 | 6 | | | 115 (2) |
| Experiment c | 118 (2) | 36 | | | | 154 (2) |
| Experiment d | 20 | 16 | 8 | 6 | | 50 |
| Experiment e | 99 | 108 (2) | 61 (1) | 8 | 2 | 278 (3) |
| Experiment f | 45 (3) | | | | | 45 (3) |
| Experiment g | 61 (3) | 37 | 4 | | | 102 (3) |
| Total | 458 (10) | 267 (2) | 103 (1) | 17 | 4 | 849 (13) |

Considering all co-cultured cells across all 7 experiments, a total of 2,363 control cells were tracked, amongst which 111 became apoptotic. The number of mitotic events is clear from the number of cells tracked in successive generations. Note that very occasionally, a daughter cell was quickly lost from vision and so not tracked, accounting for odd numbers in columns for generations 1 onwards, that contain otherwise paired sister cells.

**Table S3.** Median values for cell-profile area, cell circularity and cell migration velocity, in control SAOS-2 and HDF cultured in isolation for all experiments.

| | Median Cell-Profile Area (μm$^2$) | | Median Cell Circularity | | Median Cell Migration Velocity (μm/day) | |
|---|---|---|---|---|---|---|
| | SAOS-2 | HDF | SAOS-2 | HDF | SAOS-2 | HDF |
| Experiment a | 1,256 | 4,179 | 0.79 | 0.35 | 92 | 456 |
| Experiment b | 1,684 | 12,784 | 0.64 | 0.17 | 84 | 288 |
| Experiment c | 1,811 | 6,856 | 0.70 | 0.37 | 76 | 220 |
| Experiment d | 1,359 | 4,172 | 0.67 | 0.36 | 86 | 327 |
| Experiment e | 1,380 | 3,785 | 0.85 | 0.38 | 81 | 402 |
| Experiment f | 1,084 | 2,923 | 0.87 | 0.31 | 67 | 229 |
| Experiment g | 1,025 | 3,178 | 0.65 | 0.41 | 98 | 234 |
| Median of All Experiments | 1,359 | 4,172 | 0.70 | 0.36 | 84 | 288 |

SAOS-2 cultured in isolation had lower cell-profile area and cell migration velocity, as well as higher cell circularity, compared with HDF similarly cultured in isolation ($p < 0.016$, Wilcoxon Signed Rank Test)

**Table S4** Kendall's tau of correlation between cell-profile area of tracked SAOS-2 and absolute fluorescence acquired from co-cultured HDF (Fa), as well as with compensation for halving of fluorescence from mother cells by cell division (Fmc). Results for all experiments are shown, considering: all generations of cells together; starting and first generations together; and where a second generation of cells was present, starting, first and second generations together. Statistical significance is given, where *NS* indicates 'not significant' to $p < 0.05$. Where statistical significance was approached but not reached, the calculated p value is given (*NS (p value)*).

| | **Kendall's tau of Correlation Between Cell-Profile Area and Receipt of HDF Fluorescence by SAOS-2** | |
|---|---|---|
| | Fa | Fmc |
| **Experiment a** | | |
| Kendall's tau of all Generations Considered Together | 0.55 | 0.53 |
| *Statistical Significance of the Above* | *< 0.0001* | *< 0.0001* |
| Kendall's tau of Starting and First Generations Considered Together | 0.54 | 0.47 |
| *Statistical Significance of the Above* | *< 0.0001* | *< 0.0001* |
| **Experiment b** | | |
| Kendall's tau of all Generations Considered Together | 0.15 | 0.11 |
| *Statistical Significance of the Above* | *< 0.0001* | *< 0.0001* |
| Kendall's tau of Starting and First Generations Considered Together | 0.14 | 0.10 |
| *Statistical Significance of the Above* | *< 0.0001* | *0.0003* |
| **Experiment c** | | |
| Kendall's tau of all Generations Considered Together | 0.76 | 0.68 |
| *Statistical Significance of the Above* | *< 0.0001* | *< 0.0001* |
| Kendall's tau of Starting and First Generations Considered Together | 0.76 | 0.67 |
| *Statistical Significance of the Above* | *< 0.0001* | *< 0.0001* |
| **Experiment d** | | |
| Kendall's tau of all Generations Considered Together | 0.51 | 0.45 |
| *Statistical Significance of the Above* | *< 0.0001* | *< 0.0001* |
| Kendall's tau of Starting and First Generations Considered Together | 0.49 | 0.48 |
| *Statistical Significance of the Above* | *< 0.0001* | *< 0.0001* |
| Kendall's tau of Starting, First and Second Generations Considered Together | 0.49 | 0.44 |
| *Statistical Significance of the Above* | *< 0.0001* | *< 0.0001* |
| **Experiment e** | | |
| Kendall's tau of all Generations Considered Together | 0.13 | 0.08 |
| *Statistical Significance of the Above* | *0.001* | *0.04* |
| Kendall's tau of Starting and First Generations Considered Together | 0.09 | 0.035 |
| *Statistical Significance of the Above* | *NS (0.098)* | *NS* |
| Kendall's tau of Starting, First and Second Generations Considered Together | 0.11 | 0.07 |
| *Statistical Significance of the Above* | *0.0059* | *0.072* |
| **Experiment f** | | |
| Kendall's tau of all Generations Considered Together | 0.55 | 0.51 |
| *Statistical Significance of the Above* | *< 0.0001* | *< 0.0001* |
| Kendall's tau of Starting and First Generations Considered Together | 0.59 | 0.53 |
| *Statistical Significance of the Above* | *< 0.0001* | *< 0.0001* |
| **Experiment g** | | |
| Kendall's tau of all Generations Considered Together | 0.54 | 0.41 |
| *Statistical Significance of the Above* | *<0.0001* | *< 0.0001* |
| Kendall's tau of Starting and First Generations Considered Together | 0.5 | 0.4 |
| *Statistical Significance of the Above* | *< 0.0001* | *< 0.0001* |
| Kendall's tau of Starting, First and Second Generations Considered Together | 0.54 | 0.41 |
| *Statistical Significance of the Above* | *< 0.0001* | *< 0.0001* |

There was strong correlation of cell-profile area with HDF fluorescence. Correlation with Fa was stronger than for Fmc ($p < 0.016$, Wilcoxon Signed Rank Test). Inclusion of progressive generations of cells in calculations for tau had little effect on the strength or statistical significance of the correlation. Data indicate no heritability of the effect of fluorescence uptake past cell division, with an index of heritability of 0.

**Table S5.** Kendall's tau for correlation in differences between paired sister cells for cell-profile area and absolute fluorescence (Fa). Statistical significance *(Stat. Sig.)* is indicated to p < 0.05, and *'NS'* for 'not significant' is recorded where statistical significance was not reached.

| | First Generation Cells | | Second Generation Cells | | First and Second Generations of Cells Considered Together | |
|---|---|---|---|---|---|---|
| | Kendall's tau | *Stat. Sig.* | Kendall's tau | *Stat. Sig.* | Kendall's tau | *Stat. Sig.* |
| Experiment a | 0.31 | *0.0031* | 0.59 | *< 0.0001* | 0.41 | *0.0001* |
| Experiment b | 0.15 | *< 0.0001* | 0.12 | *NS* | 0.15 | *0.003* |
| Experiment c | 0.63 | *< 0.0001* | | | | |
| Experiment d | 0.39 | *0.011* | 0.49 | *0.0018* | 0.45 | *< 0.0001* |
| Experiment e | 0.54 | *NS* | -0.087 | *NS* | 0.02 | *NS* |
| Experiment f | 0.71 | *< 0.0001* | 0.6 | *NS* | 0.68 | *< 0.0001* |
| Experiment g | 0.36 | *< 0.0001* | 0.5 | *< 0.0001* | 0.43 | *< 0.0001* |

Divergence from an expected correlation of 0 was statistically significant as per One Sample Wilcoxon Test, for first generation cells (p < 0.016) and when first and second generation cells were considered together (p < 0.032); and approached but did not reach statistical significance for second generation cells (p = 0.063). Strong correlation was seen across experiments with exception of experiment 'e', where there was very weak negative correlation for second generation cells, and the correlation for first generation cells although strong, was not statistically significant.

**Table S6.** Kendall's tau of correlation between cell circularity of tracked SAOS-2 and absolute fluorescence acquired from co-cultured HDF (Fa), as well as with compensation for halving of fluorescence from mother cells by cell division (Fmc). Results for all experiments are shown, considering: all generations of cells together; starting and first generations together; and where a second generation of cells was present, starting, first and second generations together. Statistical significance is given, where *NS* indicates 'not significant' to p < 0.05. Where statistical significance was approached but not reached, the calculated p value is given (*NS (p value)*).

| | **Kendall's tau of Correlation Between Cell Circularity and Receipt of HDF Fluorescence by SAOS-2** | |
|---|---|---|
| | Fa | Fmc |
| **Experiment a** | | |
| Kendall's tau of all Generations Considered Together | -0.14 | -0.18 |
| *Statistical Significance of the Above* | *0.0058* | *0.0002* |
| Kendall's tau of Starting and First Generations Considered Together | -0.19 | -0.22 |
| *Statistical Significance of the Above* | *0.0009* | *< 0.0001* |
| **Experiment b** | | |
| Kendall's tau of all Generations Considered Together | -0.03 | -0.05 |
| *Statistical Significance of the Above* | *NS* | *NS (0.051)* |
| Kendall's tau of Starting and First Generations Considered Together | -0.02 | -0.05 |
| *Statistical Significance of the Above* | *NS* | *NS (0.063)* |
| **Experiment c** | | |
| Kendall's tau of all Generations Considered Together | -0.36 | -0.37 |
| *Statistical Significance of the Above* | *< 0.0001* | *<0.0001* |
| Kendall's tau of Starting and First Generations Considered Together | -0.35 | -0.36 |
| *Statistical Significance of the Above* | *< 0.0001* | *< 0.0001* |
| **Experiment d** | | |
| Kendall's tau of all Generations Considered Together | -0.11 | -0.12 |
| *Statistical Significance of the Above* | *NS (0.06)* | *0.045* |
| Kendall's tau of Starting and First Generations Considered Together | -0.24 | -0.23 |
| *Statistical Significance of the Above* | *0.0024* | *0.0032* |
| Kendall's tau of Starting, First and Second Generations Considered Together | -0.15 | -0.15 |
| *Statistical Significance of the Above* | *0.02* | *0.015* |
| **Experiment e** | | |
| Kendall's tau of all Generations Considered Together | -0.18 | 0.03 |
| *Statistical Significance of the Above* | *NS (0.14)* | *NS* |
| Kendall's tau of Starting and First Generations Considered Together | 0.10 | 0.16 |
| *Statistical Significance of the Above* | *NS (0.06)* | *0.0026* |
| Kendall's tau of Starting, First and Second Generations Considered Together | -0.04 | 0.07 |
| *Statistical Significance of the Above* | *NS* | *NS (0.096)* |
| **Experiment f** | | |
| Kendall's tau of all Generations Considered Together | -0.08 | -0.14 |
| *Statistical Significance of the Above* | *NS* | *0.019* |
| Kendall's tau of Starting and First Generations Considered Together | -0.12 | -0.16 |
| *Statistical Significance of the Above* | *0.061* | *0.012* |
| **Experiment g** | | |
| Kendall's tau of all Generations Considered Together | -0.19 | -0.15 |
| *Statistical Significance of the Above* | *< 0.0001* | *< 0.0001* |
| Kendall's tau of Starting and First Generations Considered Together | -0.22 | -0.18 |
| *Statistical Significance of the Above* | *< 0.0001* | *0.0003* |
| Kendall's tau of Starting, First and Second Generations Considered Together | -0.18 | -0.15 |
| *Statistical Significance of the Above* | *< 0.0001* | *< 0.0001* |

There was weak inverse correlation of cell circularity with HDF fluorescence in all but one experiment, where the reverse effect was seen (Exp. e). Correlations were generally strongest considering starting and first generations together, suggestive of confounding effects of cell crowding at later time points. For this reason, analysis focused on starting and first generation cells pooled. The strength of the inverse correlation across Fa and Fmc varied amongst experiments. Tau values Fmc were higher than Fa in 4 experiments (a, b, c and f), while the reverse was the case in 2 experiments (d and g), and experiment 'e' was equivocal. Although statistically compelling within a number of individual experiments, differences between experiments were such that no statistically significant result could be attributed to these general patterns. Overall, data indicate negative correlation of cell circularity with uptake of fibroblast fluorescence, with moderately strong persistence of circularity past mother cell division (index of persistence of 0.71).

**Table S7.** Kendall's tau for correlation of differences between paired sister cells for cell circularity and absolute fluorescence (Fa). Statistical significance *(Stat. Sig.)* is indicated to $p < 0.05$, and *'NS'* for 'not significant' is recorded where statistical significance was not reached. Where statistical significance was approached but not reached, calculated significance is given (*NS (p value)*).

| | First Generation Cells | | Second Generation Cells | | First and Second Generations of Cells Considered Together | |
|---|---|---|---|---|---|---|
| | Kendall's tau | *Stat. Sig.* | Kendall's tau | *Stat. Sig.* | Kendall's tau | *Stat. Sig.* |
| Experiment a | 0.06 | *NS* | -0.08 | *NS* | 0.01 | *NS* |
| Experiment b | -0.13 | *NS* | -0.04 | *NS* | -0.02 | *NS* |
| Experiment c | 0.04 | *NS* | | | | |
| Experiment d | -0.03 | *NS* | 0.18 | *NS* | 0.08 | *NS* |
| Experiment e | 0.01 | *NS* | 0.16 | *NS* | 0.12 | *0.046* |
| Experiment f | 0.60 | *NS* | 0.15 | *NS* | 0.15 | *NS* |
| Experiment g | 0.18 | *NS (0.055)* | 0.08 | *NS* | 0.14 | *0.03* |

Weak correlation was seen in only two experiments (e, g) when all cells were considered together.

**Table S8.** Kendall's tau of correlation between cell migration velocity of tracked SAOS-2 and absolute fluorescence acquired from co-cultured HDF (Fa), as well as with compensation for halving of fluorescence from mother cells by cell division (Fmc). Results for all experiments are shown, considering: all generations of cells together; starting and first generations together; and where a second generation of cells was present, starting, first and second generations together. Statistical significance is given, where *NS* indicates 'not significant' to $p < 0.05$. Where statistical significance was approached but not reached, the calculated p value is given *(NS (p value))*.

| | Kendall's tau of Correlation Between Cell Migration Velocity and Receipt of HDF Fluorescence by SAOS-2 | |
|---|---|---|
| | Fa | Fmc |
| **Experiment a** | | |
| Kendall's tau of all Generations Considered Together | 0.10 | 0.13 |
| *Statistical Significance of the Above* | *NS (0.051)* | *0.009* |
| Kendall's tau of Starting and First Generations Considered Together | 0.10 | 0.18 |
| *Statistical Significance of the Above* | *NS (0.088)* | *0.0019* |
| **Experiment b** | | |
| Kendall's tau of all Generations Considered Together | 0.03 | -0.004 |
| *Statistical Significance of the Above* | *NS* | *NS* |
| Kendall's tau of Starting and First Generations Considered Together | 0.02 | -0.02 |
| *Statistical Significance of the Above* | *NS* | *NS* |
| **Experiment c** | | |
| Kendall's tau of all Generations Considered Together | 0.24 | 0.19 |
| *Statistical Significance of the Above* | *< 0.0001* | *<0.0002* |
| Kendall's tau of Starting and First Generations Considered Together | 0.25 | 0.19 |
| *Statistical Significance of the Above* | *< 0.0001* | *< 0.0002* |
| **Experiment d** | | |
| Kendall's tau of all Generations Considered Together | 0.03 | -0.03 |
| *Statistical Significance of the Above* | *NS* | *NS* |
| Kendall's tau of Starting and First Generations Considered Together | 0.06 | 0.04 |
| *Statistical Significance of the Above* | *NS* | *NS* |
| Kendall's tau of Starting, First and Second Generations Considered Together | 0.08 | 0.01 |
| *Statistical Significance of the Above* | *0.2* | *NS* |
| **Experiment e** | | |
| Kendall's tau of all Generations Considered Together | 0.08 | 0.06 |
| *Statistical Significance of the Above* | *0.045* | *0.11* |
| Kendall's tau of Starting and First Generations Considered Together | 0.003 | -0.045 |
| *Statistical Significance of the Above* | *NS* | *NS* |
| Kendall's tau of Starting, First and Second Generations Considered Together | 0.07 | 0.04 |
| *Statistical Significance of the Above* | *NS* | *NS* |
| **Experiment f** | | |
| Kendall's tau of all Generations Considered Together | 0.19 | 0.15 |
| *Statistical Significance of the Above* | *0.0017* | *0.013* |
| Kendall's tau of Starting and First Generations Considered Together | 0.17 | 0.14 |
| *Statistical Significance of the Above* | *0.0065* | *0.025* |
| **Experiment g** | | |
| Kendall's tau of all Generations Considered Together | 0.1 | 0.011 |
| *Statistical Significance of the Above* | *0.0097* | *NS* |
| Kendall's tau of Starting and First Generations Considered Together | 0.05 | -0.04 |
| *Statistical Significance of the Above* | *NS* | *NS* |
| Kendall's tau of Starting, First and Second Generations Considered Together | 0.11 | 0.02 |
| *Statistical Significance of the Above* | *0.0069* | *NS* |

Very weak correlation of cell migration with HDF fluorescence was seen in most experiments, but this reached statistical significance in only 5 of 7 of these (a, c, e, f and g). The correlation was strongest for Fa in all experiments other than in 'a' where the strongest correlation was for Fmc. Dependent on which grouping of generations were considered, extremely weak negative correlations were seen for Fmc in four experiments (b, d, e, f), and none of these were statistically significant, so it seems reasonable to reject them. There was no statistically significant difference between Fa and Fmc. Comparison of Fa with Fmc provided negligible evidence for any persistence of the effect beyond cell division, with a calculated index of 0.14. Overall, data suggest a modest association between fluorescence uptake and cell migration, that is not persistent past cell division.

**Table S9.** Kendall's tau for correlation of differences between paired sister cells for cell migration velocity and absolute fluorescence (Fa). Statistical significance *(Stat. Sig.)* is indicated to $p < 0.05$, and *'NS'* for 'not significant' is recorded where statistical significance was not reached.

| | First Generation Cells | | Second Generation Cells | | First and Second Generations of Cells Considered Together | |
|---|---|---|---|---|---|---|
| | Kendall's tau | *Stat. Sig.* | Kendall's tau | *Stat. Sig.* | Kendall's tau | *Stat. Sig.* |
| Experiment a | -0.11 | *NS* | -0.02 | *NS* | -0.11 | *NS* |
| Experiment b | -0.05 | *NS* | -0.02 | *NS* | -0.06 | *NS* |
| Experiment c | 0.15 | *NS* | | | | |
| Experiment d | -0.12 | *NS* | 0.23 | *NS* | -0.04 | *NS* |
| Experiment e | 0.01 | *NS* | 0.16 | *NS* | 0.10 | *NS* |
| Experiment f | 0.14 | *NS* | 0.60 | *NS* | 0.16 | *NS* |
| Experiment g | 0.09 | *NS* | 0.10 | *NS* | 0.12 | *NS* |

No convincing statistically significant correlations were seen.
